# ALOXE3 is a hepatic fasting-responsive lipoxygenase that enhances insulin sensitivity via hepatic PPARγ

**DOI:** 10.1101/267781

**Authors:** Cassandra B. Higgins, Yiming Zhang, Allyson L. Mayer, Hideji Fujiwara, Alicyn I. Stothard, Mark J. Graham, Benjamin M. Swarts, Brian J. DeBosch

**Affiliations:** Department of Pediatrics, Washington University School of Medicine, St. Louis, MO 63110; Department of Medicine, Diabetic Cardiovascular Disease Center, Washington University School of Medicine, St. Louis, MO 63110; Department of Chemistry & Biochemistry, Central Michigan University, Mt. Pleasant, MI 48859; IONIS Pharmaceuticals and 6Cell Biology and Physiology, Washington University School of Medicine, St. Louis, MO 63110; Department of Cell Biology & Physiology, Washington University School of Medicine, St. Louis, MO 63110

**Keywords:** **Liver**, **energy metabolism**, **insulin resistance**, **glucose transport**, **non-alcoholic fatty liver disease**, **obesity**, **PPARγ**, **thiazolidinediones**, **peroxisome proliferator antigen receptor-gamma co-receptor-1α**, **lipoxygenase**, **lipids**, **ALOXE3**, **trehalose**, **lactotrehalose**

## Abstract

The hepatic glucose fasting response is gaining traction as a therapeutic pathway to enhance hepatic and whole-host metabolism. However, the mechanisms underlying these metabolic effects remain unclear. Here, we demonstrate the lipoxygenase, ALOXE3, is a novel effector of the thepatic fasting response. We show that ALOXE3 is activated during fasting, glucose withdrawal, and trehalose/trehalose analogue treatment. Hepatocyte-specific ALOXE3 expression reduced weight gain and hepatic steatosis in dietaryand genetically obese (db/db) models. ALOXE3 expression moreover enhanced basal thermogenesis and abrogated insulin resistance in db/db diabetic mice. Targeted metabolomics demonstrated accumulation of the PPARγ ligand, 12-KETE in hepatocytes overexpressing ALOXE3. Strikingly, PPARγ inhibition reversed hepatic ALOXE3-mediated insulin sensitization, suppression of hepatocellular ATP production and oxygen consumption, and gene induction of PPARγ coactivator-1a (PGC1α) expression. Moreover, hepatocyte-specific PPARγ deletion reversed the therapeutic effect of hepatic ALOXE3 expression on diet-induced insulin intolerance. ALOXE3 is therefore a novel effector of the hepatocellular fasting response that leverages both PPARγ-mediated and pleiotropic effects to augment hepatic and whole-host metabolism, and is thus a promising target to ameliorate metabolic disease.

## INTRODUCTION

Leveraging the generalized fasting and caloric restriction responses to mitigate metabolic disease is under intense investigation as a novel therapeutic approach against obesity, diabetes mellitus and non-alcoholic fatty liver disease (NAFLD) (1,2). The hepatocyte is uniquely positioned at the nexus of the portal circulation - where dietary macronutrients are sensed and triaged to their metabolic fate - and the peripheral circulation, from which macronutrients are taken up and utilized or stored. Its unique anatomic positioning also defines the hepatocyte’s role in coordinating the transition from fed to fasting states. Specifically, the hepatocyte senses fat, protein and carbohydrate content and rapidly alters its metabolism and its endocrine functions to coordinate peripheral responses. However, the promise of caloric restriction and generalized macronutrient fasting remains limited by its clinical impracticality and unsustainability. Our laboratory and others have therefore examined specific pathways through which fasting and fasting-like mechanisms can be molecularly or pharmacologically leveraged to the benefit of the host.

Upon physiological macronutrient withdrawal, the canonical hepatic response is to shift to mobilize glycogen and oxidize fat from stores to produce glucose and ketone body fuel for the brain and heart. This begins with activation of proximal macronutrient “sensing” kinases, AMP-activated protein kinase (AMPK) and the mechanistic target of rapamycin (mTOR). These kinases act as primary regulators of macroautophagy as well as transcriptional fasting programming by hepatic transcription factors, including transcription factor EB (3), PPARγ coactivator-1 (4) and transcriptional modifiers such as the sirtuin family of protein deacetylases (5). FGF21 is released from the liver of fasting rodents and humans to promote adipose “browning” (mitochondrial uncoupling to promote heat generation (6)). The net effect of this signaling initiation is a networked cascade through which the liver provides energy substrate to the periphery during fasting, and promotes peripheral insulin sensitivity for efficient absorption of the next meal.

We recently demonstrated that blocking hepatic glucose transporters genetically or pharmacologically closely recapitulates the therapeutic metabolic effects of generalized macronutrient fasting (7–9). Pharmacologic glucose transport blockade using the disaccharide glucose transporter inhibitor, trehalose, blocked mTOR and activated AMPK-dependent autophagic flux (10,11). Moreover, trehalose blocked diet- and genetically induced steatosis via ATG16L1- and AMPK-dependent mechanisms, and enhanced peripheral thermogenesis via hepatocyte TFEB- and FGF21-dependent mechanisms (data in revision for publication). Consistent with data from pharmacological GLUT blockade, genetic deletion of the hepatocyte glucose transporter (GLUT), GLUT8, similarly increased basal heat generation, enhanced hepatocyte fat oxidation and inhibited hepatic de novo lipogenesis and triacylglycerol generation in mice fed a high-fructose diet (7,12). Notably, in those experiments, GLUT blockade was associated with enhanced enhanced insulin tolerance, enhanced hepatic PPARγ expression and concomitant activation of the PPARγ target, UCP2 (12). Conversely, GLUT8 was upregulated in the livers of genetically diabetic mice (13). Together, prior data suggest that pharmacologic and genetic activation of hepatic glucose fasting-like responses are sufficient to mitigate metabolic disease. However, the mechanisms underlying this protection remain enigmatic.

Recent data illuminate a role for cellular lipoxygenases in the pathogenesis and protection from metabolic disease. In humans, a recent large cohort adults with type 2 diabetes demonstrated a potentially protective role for 5-lipoxygenases ALOX5 and ALOX5AP against diabetes mellitus (14). Conversely, ALOX12/15 inhibition in rodent models inhibited oxidative stress, β-cell deterioration, hepatic steatosis and systemic inflammation (15‒18). Together, these data suggest that intracellular lipid intermediary metabolism drives both tissue-level and whole-organism metabolic homeostasis. The concept that the fasting state might modulate lipoxygenase activity as part of the broadly adaptive effects of fasting on metabolism remained heretofore unexplored.

Here, we identified ALOXE3 as a fasting-induced epidermal-type lipoxygenase that acts as a novel effector of the hepatic glucose fasting response *in vitro* and *in vivo*. Targeted ALOXE3 overexpression induced robust generation of the PPARγ ligand (19,20), 12-KETE. In vivo, hepatic ALOXE3 expression enhanced insulin sensitivity, attenuated weight gain and reduced hepatic steatosis in both diet- and genetically induced steatosis models. Moreover, hepatic ALOXE3 expression mitigated diet-induced dyslipidemia, and db/db mice expressing ALOXE3 exhibited attenuated weight gain, enhanced basal caloric expenditure and PPARγ-dependent insulin sensitivity. Strikingly, we demonstrate that hepatocyte-specific PPARγ deletion is absolutely required for the insulin-sensitizing effect of hepatic ALOXE3 induction. Taken together, ALOXE3 is a fasting-responsive hepatocyte effector that is sufficient to attenuate insulin resistance, weight gain and hepatic fat deposition in dietary and genetically obese models, in part by activating hepatic PPARγ. ALOXE3 is thus a novel therapeutic mechanism downstream of the fasting response, which can be leveraged against metabolic disease.

## RESULTS

To test the hypothesis that ALOXE3 is induced by acute fasting or pseudo-fasting conditions, we subjected WT mice to 48h fasting. During acute fasting, hepatic ALOXE3 expression was increased and sustained at least 48hrs (Fig. 1A). Because we previously demonstrated a hyper-responsive hepatic fasting response in mice lacking the hepatic glucose and fructose transporter, GLUT8, we fasted GLUT8-deficient mice under the same conditions, and observed trends toward increased hepatic ALOXE3 expression (Fig. 1A). Oral trehalose (3% PO, ad libitum), similarly induced ALOXE3 expression after 24 hrs and 48hrs trehalose feeding (Fig. 1B). We then modeled ALOXE3 expression in vitro to determine whether fasting-induced ALOXE3 expression is cell-autonomously regulated. Isolated primary murine hepatocytes were subjected to treatment by either low serum and glucose (0.5% serum, 1g/L glucose), quercetin (a flavone-class glucose transporter inhibitor) or the disaccharide GLUT substrate/inhibitor (10,11,21), trehalose (Fig. 1C). In addition, we characterized fasting-mimetic effects of a trehalose analogue that resists enzymatic degradation by trehalases (22–24), lactotrehalose (α-D-glucopyronosyl-(1,1)-α-D-galactopyranoside, Fig. 1C and Supplemental Fig. 1). In each case, ALOXE3 expression was significantly increased, most potently by trehalose and lactotrehalose (Fig. 1C and Supplemental Fig. 1).

**Figure 1.**
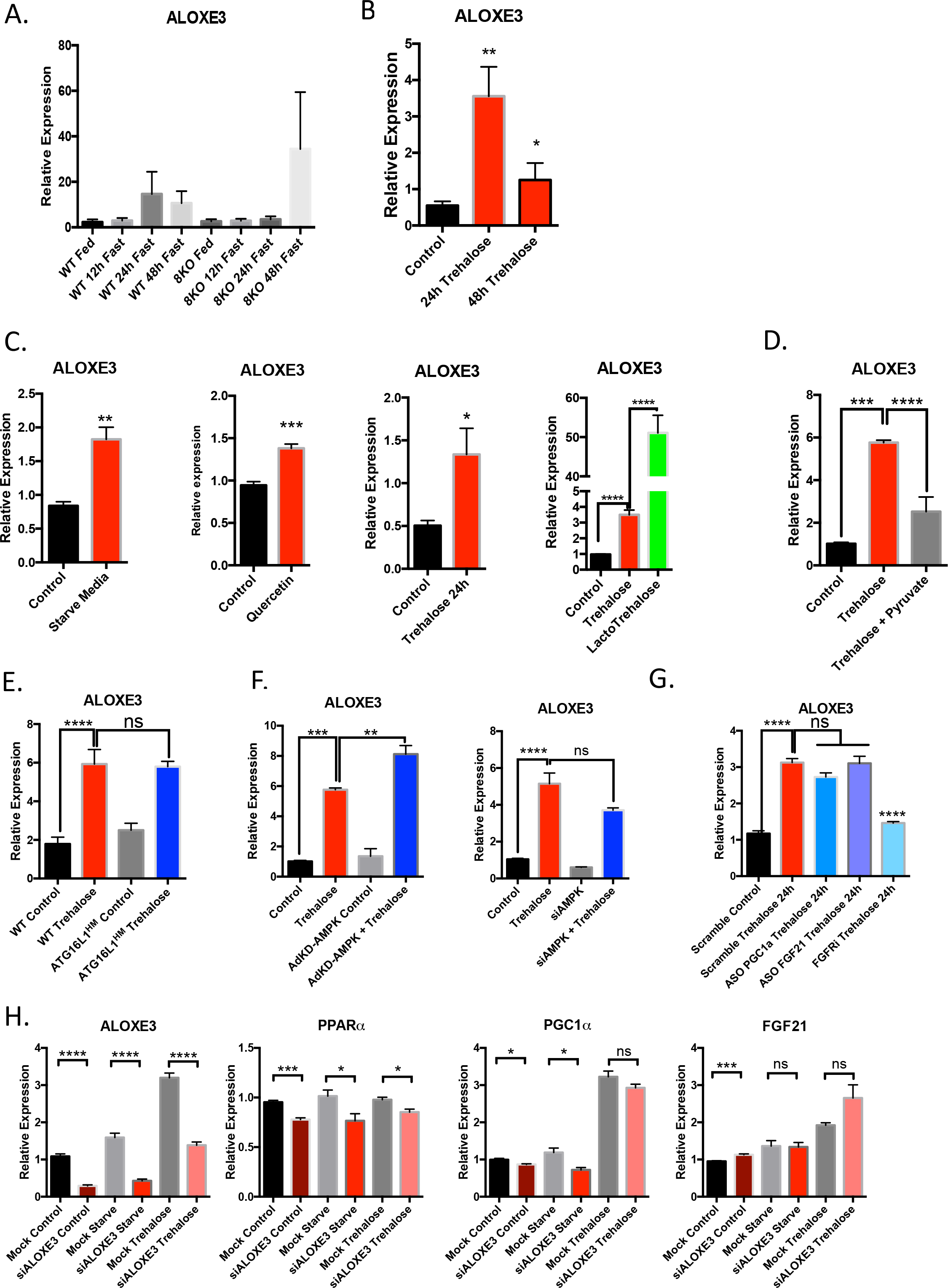
ALOXE3 is induced in response to fasting and pseudofasting. A. ALOXE3 expression in fasting WT and GLUT8-deficient mice (0-48h). B. ALOXE3 expression in WT mice fed oral trehalose (3% in sterile water fed ad libitum, 0-48h). C. ALOXE3 expression in isolated primary murine hepatocytes treated with 0.5% FCS/1g*L^−1^ glucose media, 5 μM quercetin, 100mM trehalose or 100mM lactotrehalose (24h). D. ALOXE3 expression in trehalose-treated isolated primary hepatocytes pre-treated with or without 5mM pyruvate. E. ALOXE3 expression in fasting WT and ATG16L1-mutant mice in vivo (24h). F. ALOXE3 expression in response to trehalose in the presence or absence of kinase-dead AMPK overexpression or siRNA-mediated AMPK knockdown. G. ALOXE3 expression in response to trehalose in the presence or absence of antisense oligonucleotide (ASO) directed against PPARα, PGC1α or FGF21. FGF receptor 1-4 inhibitor (LY2874455) was included as a control to demonstrate ALOXE3 blockade in context. n = 4-6 independent cultures per experiment.

To determine whether ALOXE3 induction depended upon energy substrate deficit, we treated primary hepatocytes with trehalose in the presence or absence of pyruvate. This provided energy substrate for the cell independent of glucose transporter blockade. Pyruvate suppressed trehalose-induced ALOXE3 induction by ~60% (P < 0.0001), suggesting that ALOXE3 expression is at least partly energy substrate-dependent (Fig. 1D). In light of our prior data in which we demonstrated that the autophagy complex protein ATG16L1 was required for autophagic and anti-steatotic effects of trehalose in hepatocytes (10,21), we examined whether ATG16L1 is required for ALOXE3 induction. We treated primary hepatocytes from WT littermates or from mice with homozygous hypomorphic ATG16L1 alleles (ATG16L1^HM/HM^) with or without trehalose. ALOXE3 mRNA quantification revealed robust ALOXE3 induction in wild-type hepatocytes, which was not suppressed in ATG16L1^HM/HM^ hepatocytes (Fig. 1E). Similarly, AMPK inhibition (by kinase-dead AMPK overexpression or by ALOXE3 siRNA transfection, (Fig. 1F)) and PPARα, PGC1α and FGF21 knockdown each failed to suppress ALOXE3 induction (Fig. 1G). Accordingly, moderate hepatocyte-specific overexpression of the fasting-induced transcription factor, SIRT1 did not induce ALOXE3 expression (Supp. Fig 2). In contrast, siRNA-based ALOXE3 knockdown attenuated PPARα and PGC1a at baseline and during glucose and serum withdrawal from hepatocytes in vitro, without effect on FGF21 expression. Together, our data demonstrate that ALOXE3 is a lipoxygenase that is induced in response to hepatocyte glucose transporter blockade and energy deficit via a mechanism that does not require canonical ATG16L1-dependent and AMPK-PGC1α/PPARα-FGF21 fasting mechanisms.

ALOXE3 generates the PPARγ ligand, 12-KETE, and an epoxyalcohol from 12-HpETE in the metabolism of plasma membrane arachidonic acid ((25–29), Fig 2A). We tested whether ALOXE3 similarly mediated lipid metabolism in hepatocytes upon either forced ALOXE3 expression. ALOXE3 overexpression phenocopied trehalose treatment in primary hepatocytes, and significantly enhanced 12-KETE production (Fig. 2A). Moreover, ALOXE3 overexpression suppressed production of competing metabolic pathway products 5-HETE and 12-HETE (Fig. 2C).

**Figure 2.**
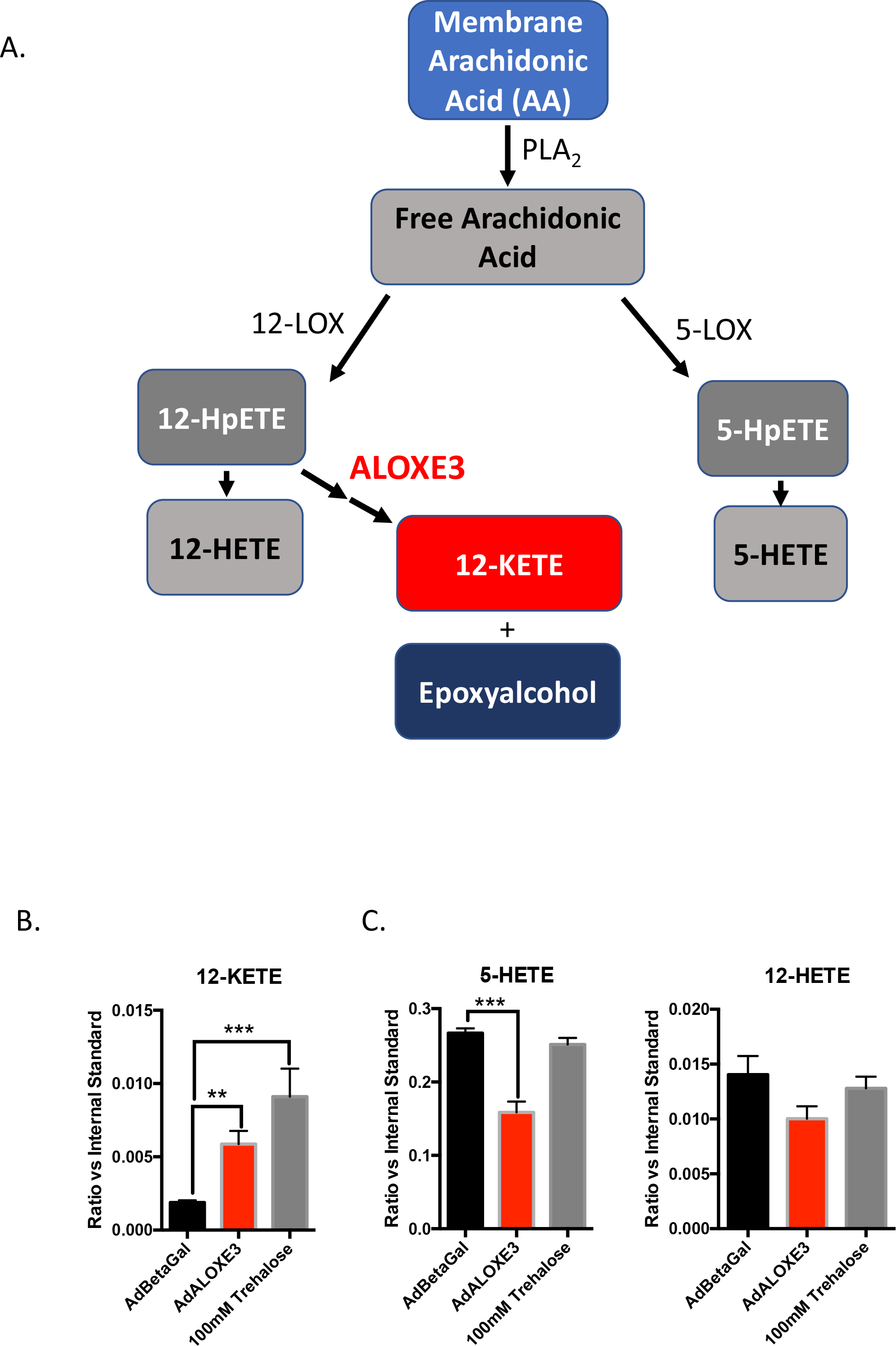
ALOXE3 and trehalose induce the PPARγ ligand 12-KETE in murine hepatocytes. A. Arachidonic acid metabolism mediated by lipoxygenases 12-LOX, 5-LOX and ALOXE3 (adapted from (26,29,58,59). B. and C. Quantitative GC-MS analysis of the stable lipoxygenase reaction products 5-HETE, 12-HETE and 12-KETE. n = 6-9 independent cultures per experiment.

To ascertain transcriptome-wide effects of forced ALOXE3 overexpression, we next treated primary hepatocytes with adenovirus encoding β-galactosidase or ALOXE3 prior to RNAseq analysis. Pathway analysis revealed that five of the ten most downregulated processes in ALOXE3-overexpressing cells were devoted to inflammatory signaling, including tumor necrosis factor-α, NF-kappa B, chemokine signaling, cytokine receptor signaling and MAPK signaling (Fig. 3A). Given these findings, and in light of the fact that ALOXE3 upregulation correlates with reduced diet- and genetically-induced steatosis (9,21) we examined whether ALOXE3 attenuates fat-induced inflammatory signaling and triglyceride accumulation. Primary hepatocytes treated with BSA-conjugated fatty acids induced IL-1β and TNFα gene expression as well as TG accumulation (Fig. 3B and 3C). Both TG accumulation and IL-1β and TNFα were reversed in FA-treated cultures overexpressing ALOXE3. The ability of ALOXE3 to attenuate FA-induced hepatic TG accumulation was partly reversed by pre-treating cultures with the lipoxygenase inhibitor, baicalein (Fig. 3D) (28).

**Figure 3.**
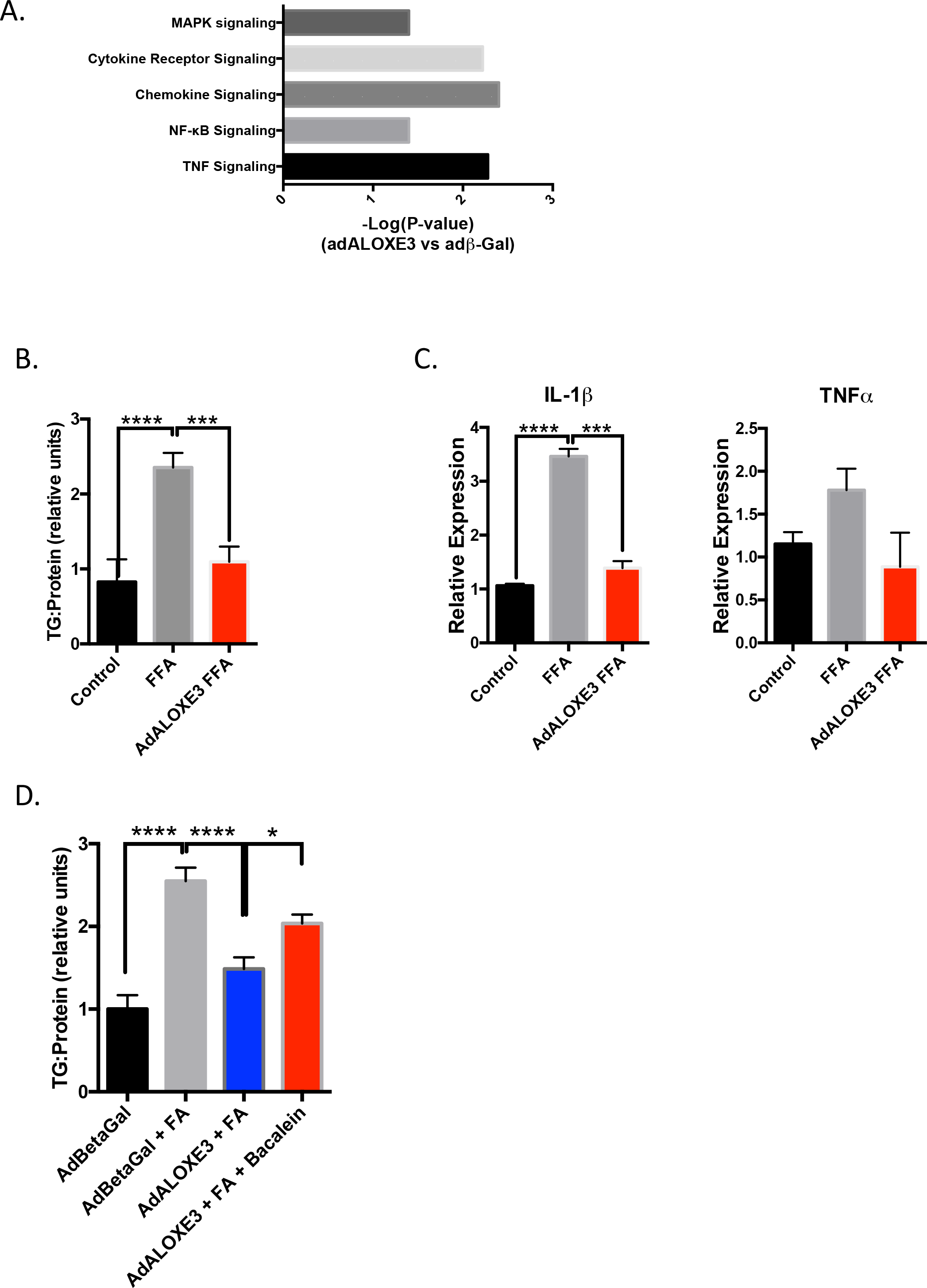
ALOXE3 reduces inflammatory signaling and steatosis in hepatocytes. A. RNAseq analysis of the top 9 down-regulated genes in primary hepatocytes expressing either beta galactosidase or ALOXE3. The P value for significantly downregulated pathways is demonstrated as (−Log(P-value)). B. Hepatic TG accumulation and C. IL-1β or TNFα expression in hepatocytes treated with BSA-conjugated fatty acids with or without ALOXE3 expression. D. Hepatic TG accumulation in hepatocytes treated with BSA-conjugated fatty acids with or without ALOXE3 expression after treatment with or without the lipoxygenase inhibitor baicalein (100 μM). n = 4-6 independent cultures per experiment.

ALOXE3-deficient mice are not amenable to in vivo metabolic studies because germline ALOXE3 deficiency results in post-natal mortality secondary to massive skin permeability and water loss. To evaluate the in vivo metabolic consequences of hepatocyte ALOXE3 activation, we tested the in vivo effects of hepatocyte-directed ALOXE3 expression. We first confirmed overexpression of the ALOXE3 transgene in liver (Fig. 4A) without changes in ALOXE3 expression in skeletal muscle, white adipose tissue or brown adipose tissue (not shown). This correlated with upregulation of fasting and oxidative genes in unperturbed ALOXE3 overexpressing mice, including ALOXE3 (Fig. 4A), PPARγ-coactivator-1α (PGC1α), hepatocyte nuclear factor 4α (HNF4α) and phosphoenolpyruvate carboxykinase 1 (PCK1, Fig 4A.). We next evaluated the effect of hepatic ALOXE3 expression in mice fed low-fat diet (LFD) or steatogenic, high trans-fat / cholesterol diet (HTFC) (12 weeks).

**Figure 4.**
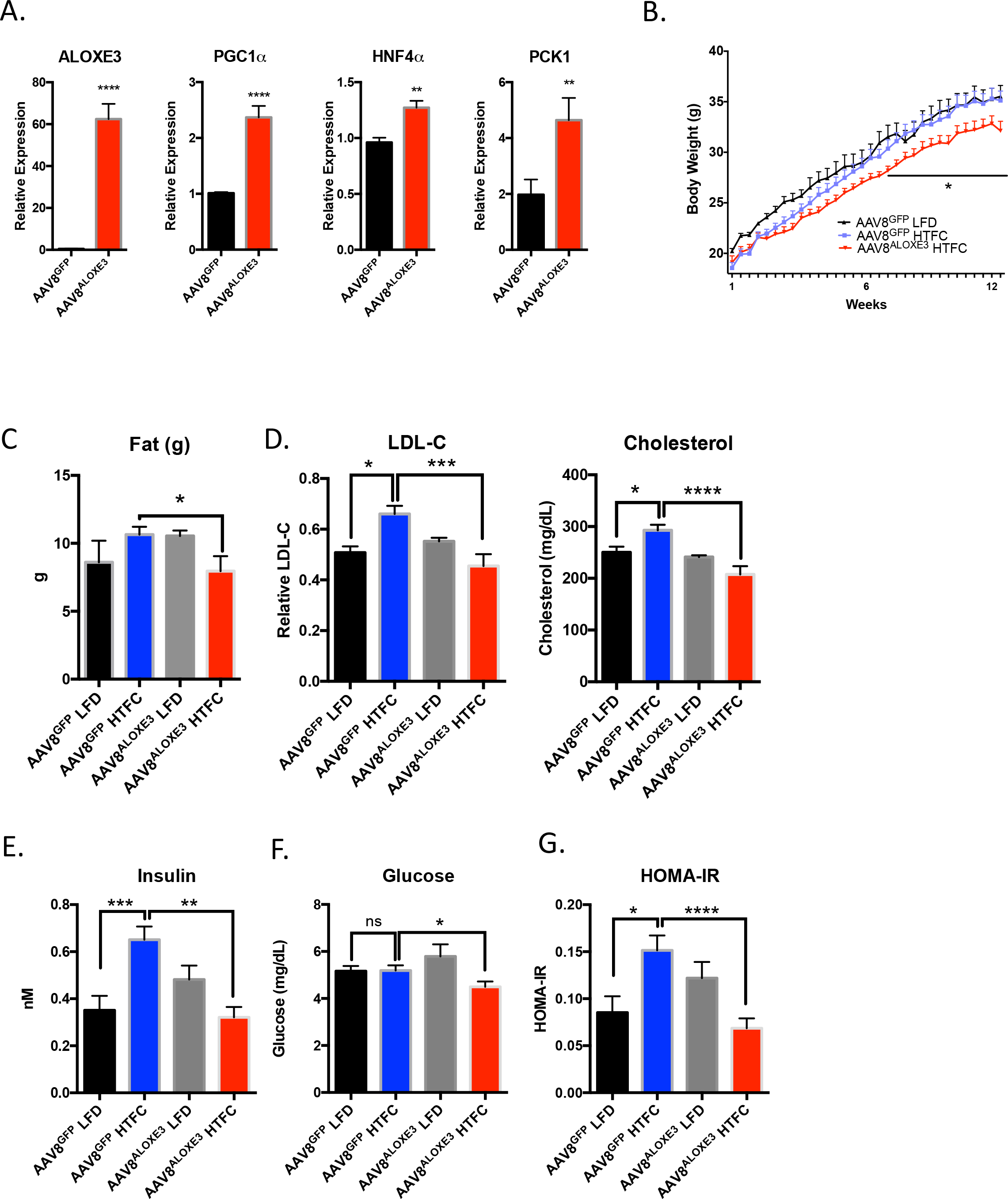
Enhanced whole-body metabolism in mice ALOXE3-overexpressing mice. A. qRT-PCR quantification of expression for oxidative and fasting-response genes in unperturbed mice expressing hepatocyte GFP or ALOXE3. B. Body weight over time in low-fat or high trans-fat/cholesterol diet-fed mice expressing hepatocyte GFP or ALOXE3. C. Body fat content in mice fed HTFC or LFD with or without hepatic ALOXE3 overexpression. D. LDL-C and total cholesterol in mice fed HTFC or LFD with or without hepatic ALOXE3 overexpression. E-G. Circulating insulin, glucose and calculated HOMA-IR in LFD- and HTFC-fed mice overexpressing hepatocyte GFP or ALOXE3. n = 5-10 mice per experiment.

Mice overexpressing ALOXE3 fed HTFC gained significantly less weight, had lower total body mass (Fig. 4B) and lower body fat mass (Fig. 5C) without changes in lean mass (not shown) when compared with GFP-overexpressing mice fed HTFC. Accordingly, LDL-C and total cholesterol were significantly lowered in HTFC-fed mice overexpressing ALOXE3 (Fig. 5D). Indices of glucose homeostasis were also improved by ALOXE3 overexpression in HTFC-fed mice (Figs. 5E, 5F, 5G). Specifically, HTFC feeding increased circulating insulin and HOMA-IR in GFP-overexpressing mice (Fig. 5E), without altering fasting plasma glucose (Fig. 5F). In contrast, HTFC-fed mice overexpressing hepatic ALOXE3 were protected from HTFC-induced hyperinsulinemia and insulin resistance (Fig. 5E and 5G). Indeed, fasting glucose was also lowered in ALOXE3-overexpressing, HTFC-fed mice when compared with GFP-expressing HTFC-fed mice (Fig. 5F). Together, hepatocyte ALOXE3 expression was sufficient to reduce diet-induced weight gain, body fat accumulation, dyslipidemia and insulin resistance.

**Figure 5.**
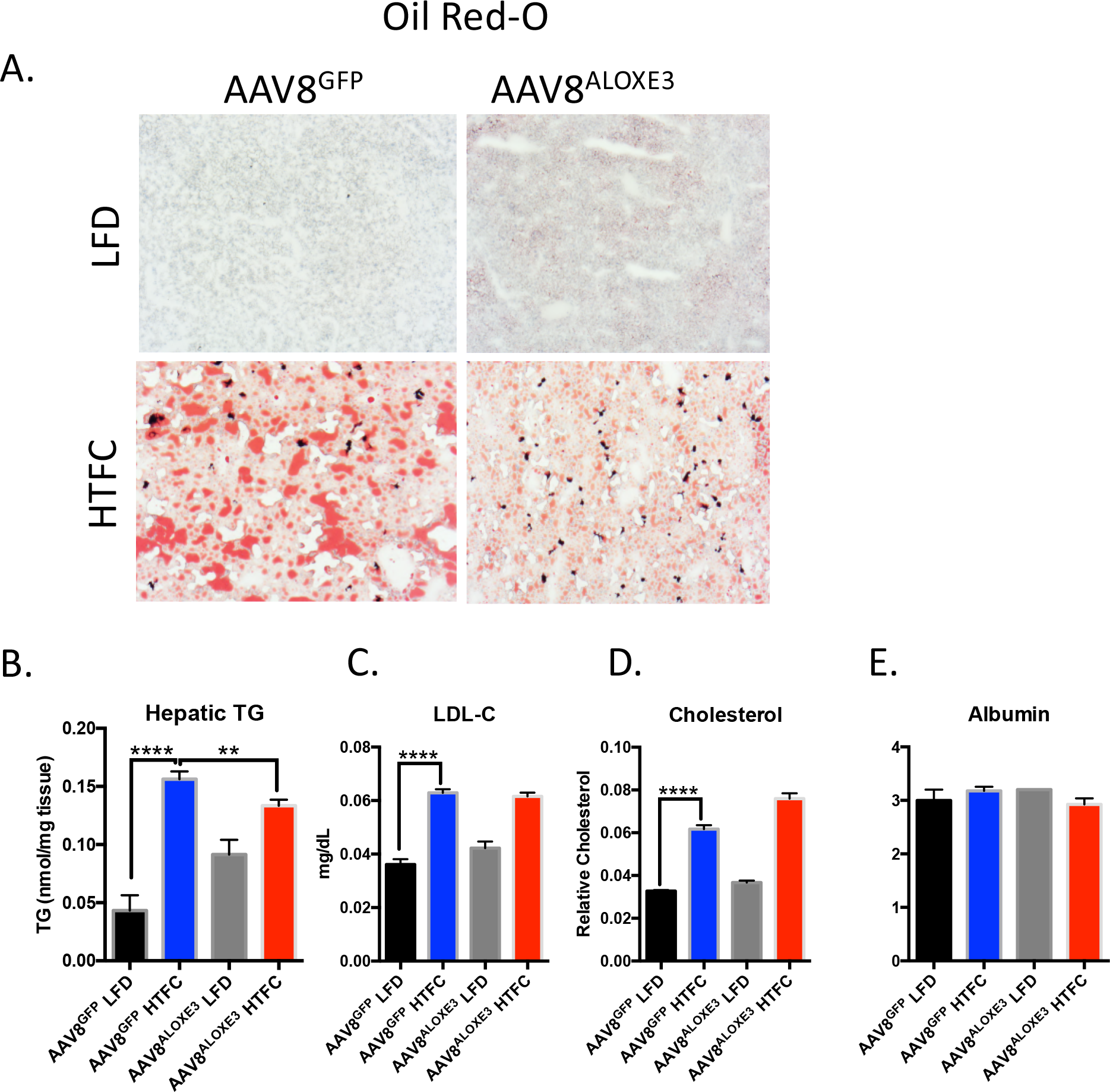
Reduced hepatic steatosis correlates with ALOXE3-induced fasting. A. Oil-red O staining in livers from low-fat- or high trans-fat/cholesterol-fed mice overexpressing empty vector or ALOXE3. B-D. Hepatic tissue quantification of triglycerides, LDL-C, total cholesterol in low-fat- or high trans-fat/cholesterol-fed mice overexpressing empty vector or ALOXE3. E. Serum albumin measurements in mice analyzed in A-D. n = 5-10 mice per experiment.

We next examined hepatic lipid metabolism in GFP- and ALOXE3-overexpressing mice fed LFD or HTFC. Frozen liver sections from HTFC-fed mice exhibited increased oil red-O staining and macrosteatosis when compared with LFD-fed mice. Consistent with prior models of PPARγ activation (30–32), ALOXE3 expression in low-fat diet-fed mice resulted in a mild basal TG accumulation (Fig. 5A and 5B). Also consistent with prior reports on PPARγ activation, ALOXE3-overexpression modestly but significantly protected from HTFC-induced TG accumulation that took on a microsteatotic staining pattern (Fig. 5A and 5B). No changes in hepatic LDL-C or total cholesterol were observed (Fig. 5C and 5D) upon hepatic ALOXE3 overexpression in LFD- or HTFC-fed mice. None of our genetic or dietary manipulations had any effect on hepatic synthetic function, as ascertained by quantitative circulating albumin levels (Fig. 5E).

To examine mechanistically how ALOXE might activate hepatocyte-starvation-like responses, we evaluated the effects of ALOXE overexpression on mitochondrial respiratory function at the cellular and molecular levels. RNAseq analysis of primary hepatocytes overexpressing ALOXE3 revealed that six of the ten most down-regulated molecular processes encompassed mitochondrial electron transport function, including proton transport, ATPase activity, transmembrane ion transport, and hydrogen export (Fig. 6A). We therefore tested functionally the hypothesis that ALOXE3 mitigates ATP production by inducing hepatocyte mitochondrial uncoupling. Seahorse analysis of hepatocytes overexpressing ALOXE3 exhibited elevated proton leak, ATP production and coupling efficiency, concomitant with suppressed basal oxygen consumption rate and enhanced glycolytic rate when compared with hepatocytes expressing beta galactosidase (Fig. 6B). Each of these parameters was partly or fully reversed in the presence of the PPARγ inhibitor, GW9662 (Fig. 6B). No changes in non-mitochondrial oxygen consumption were observed (Fig. 6C), suggesting that ALOXE3 specifically affected mitochondrial energy metabolism.

**Figure 6.**
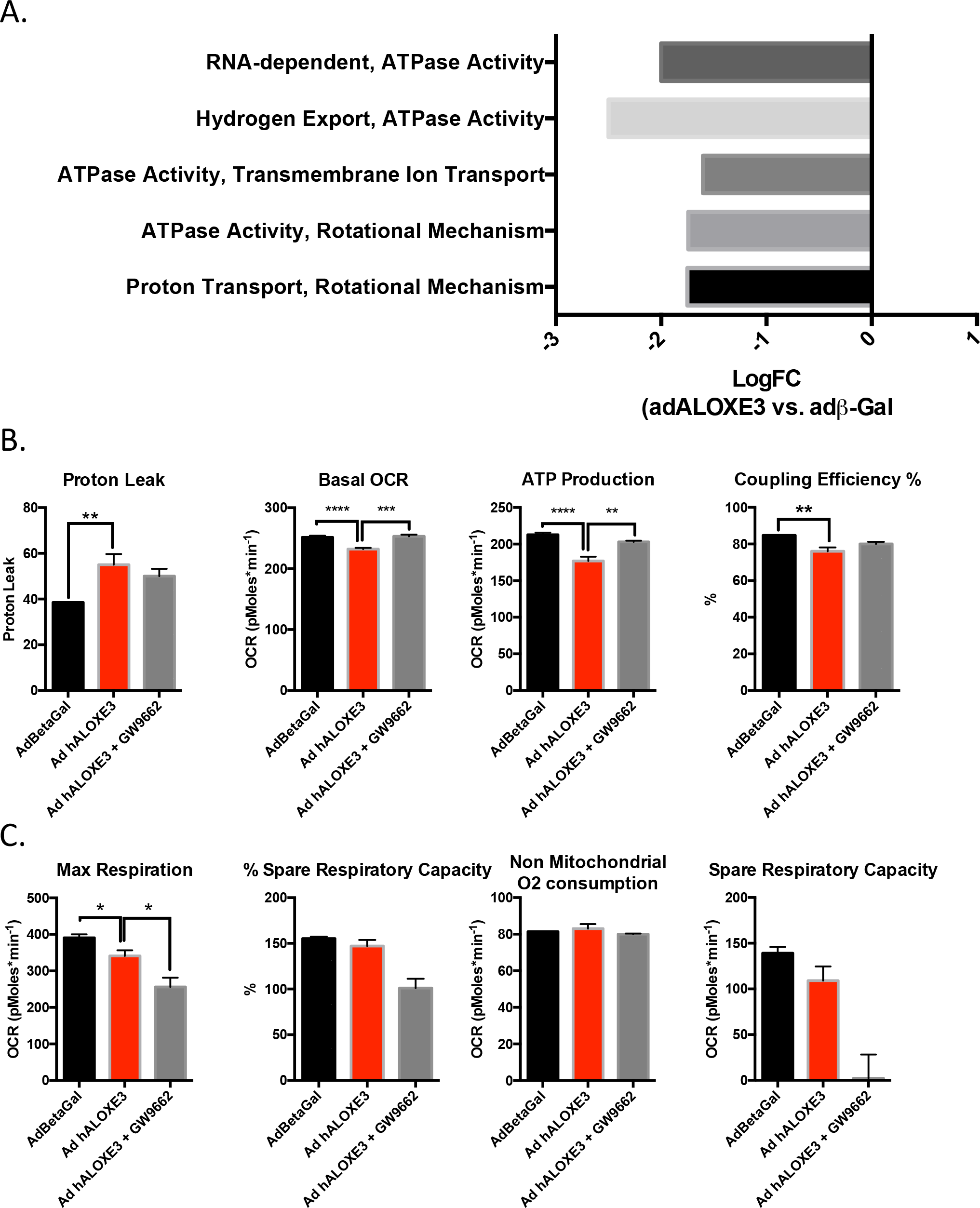
Hepatic PPARγ is required for ALOXE3 to induce hepatic energetic inefficiency. A. RNAseq analysis of the top 10 downregulated molecular processes in hepatocytes overexpressing ALOXE3. Highlighted in red are ATPase-related and mitochondrial coupling processes. Graphed is logFC relative to cultures expressing β-galactosidase. B. and C. Seahorse XF96 analysis of proton leak, basal OCR, ATP production, coupling efficiency and non-mitochondrial oxygen consumption in AML12 cells overexpressing ALOXE3 with or without GW9662 (PPARγ inhibitor) treatment. n = 6-8 independent cultures per experiment.

Mechanistic interrogation in vivo was executed to determine whether hepatocyte ALOXE3 mediated peripheral insulin and glucose homeostasis in a PPARγ-depdnent manner. To that end, we overexpressed ALOXE3 or GFP in db/db mice, which lack the leptin receptor. Again, in the liver, we observed modest, statistically significant attenuation of hepatic triglyceride accumulation in db/db mice overexpressing ALOXE3 when compared with db/db controls (Fig. 7A and 7B). Again, we confirmed ALOXE3 overexpression correlated with increases in oxidative and fasting-response genes, PGC1α, HNF4α and PCK1 (Fig. 7C). Moreover, ALOXE3 reduced genetic markers of de novo lipogenesis in db/db mice, including FSP27, SCD1 and FASN (Fig. 7D). Concomitant PPARγ inhibition by the inhibitor GW9662, statically significantly or trends toward reduction in each of these ALOXE3-induced marker genes (Fig. 7C and 7D)

**Figure 7.**
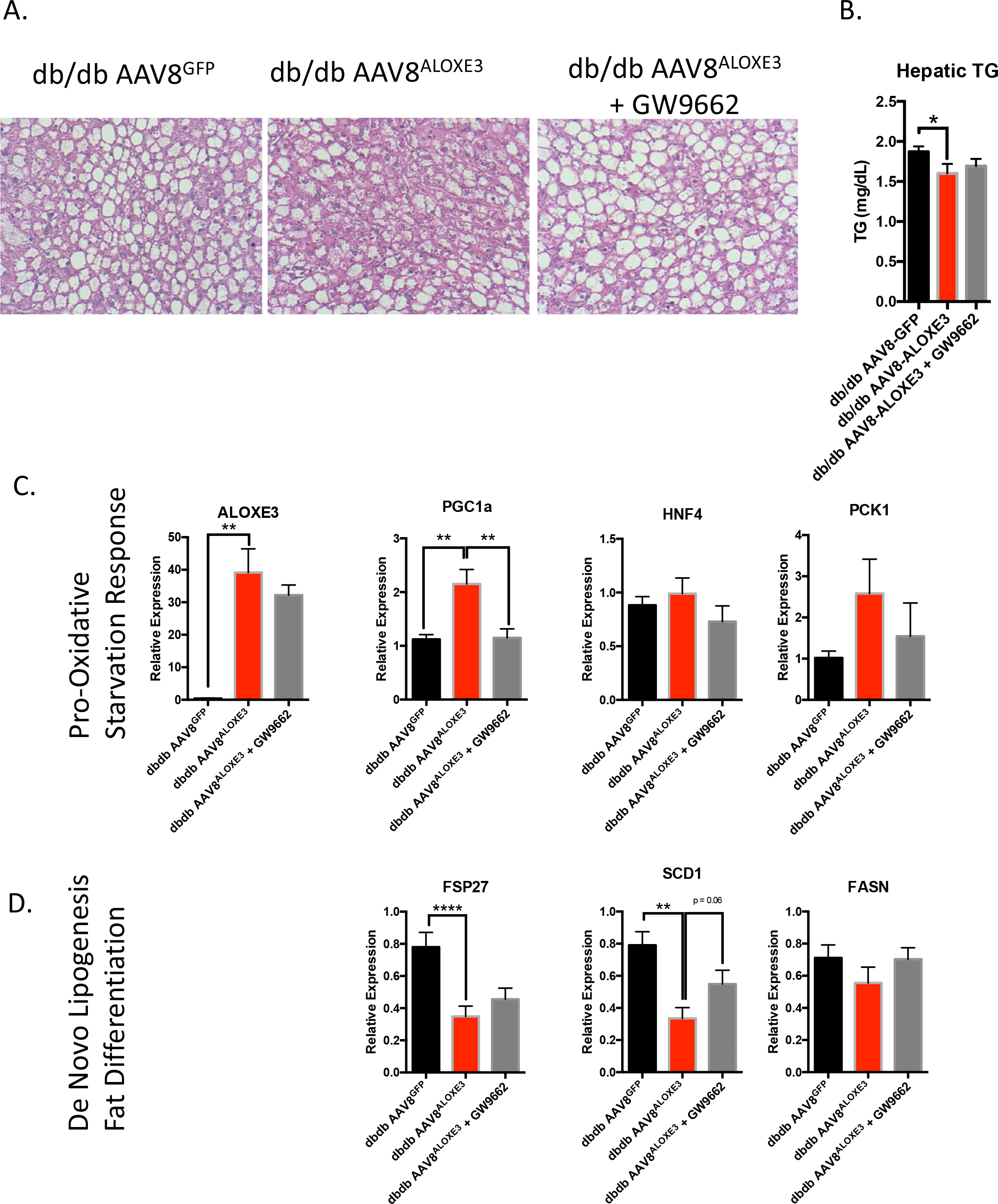
Reduced hepatic steatosis correlates with ALOXE3-induced fasting and reduction of de novo lipogenesis. A. H&E staining in livers from low-fat- or high trans-fat/cholesterol-fed mice overexpressing empty vector or ALOXE3. B. Hepatic tissue triglyceride quantification in db/db mice overexpressing GFP or ALOXE3 in the presence or absence of GW9662. C. and D. qRT-PCR quantification of expression for oxidative and fasting-response genes (C.) and de novo lipogenic genes (D.) in db/db mice expressing GFP or ALOXE3 with or without GW9662 administration. n = 5-10 mice per experiment.

Although baseline weights were not statistically different, db/db mice overexpressing ALOXE3 gained significantly less weight and had a lower end-of-trial weight than db/db mice over the 28-day trial (Fig. 8A). Indirect calorimetry revealed that the attenuated weight gain in ALOXE3^db/db^ mice was associated with enhanced light- and dark-cycle heat generation and O_2_-CO_2_ exchange (Fig. 8B and 8C). Neither heat generation nor O_2_-CO_2_ exchange in ALOXE3^db/db^ was affected by GW9662 co-administration (Fig. 8B, 8C, and 8D).

**Figure 8.**
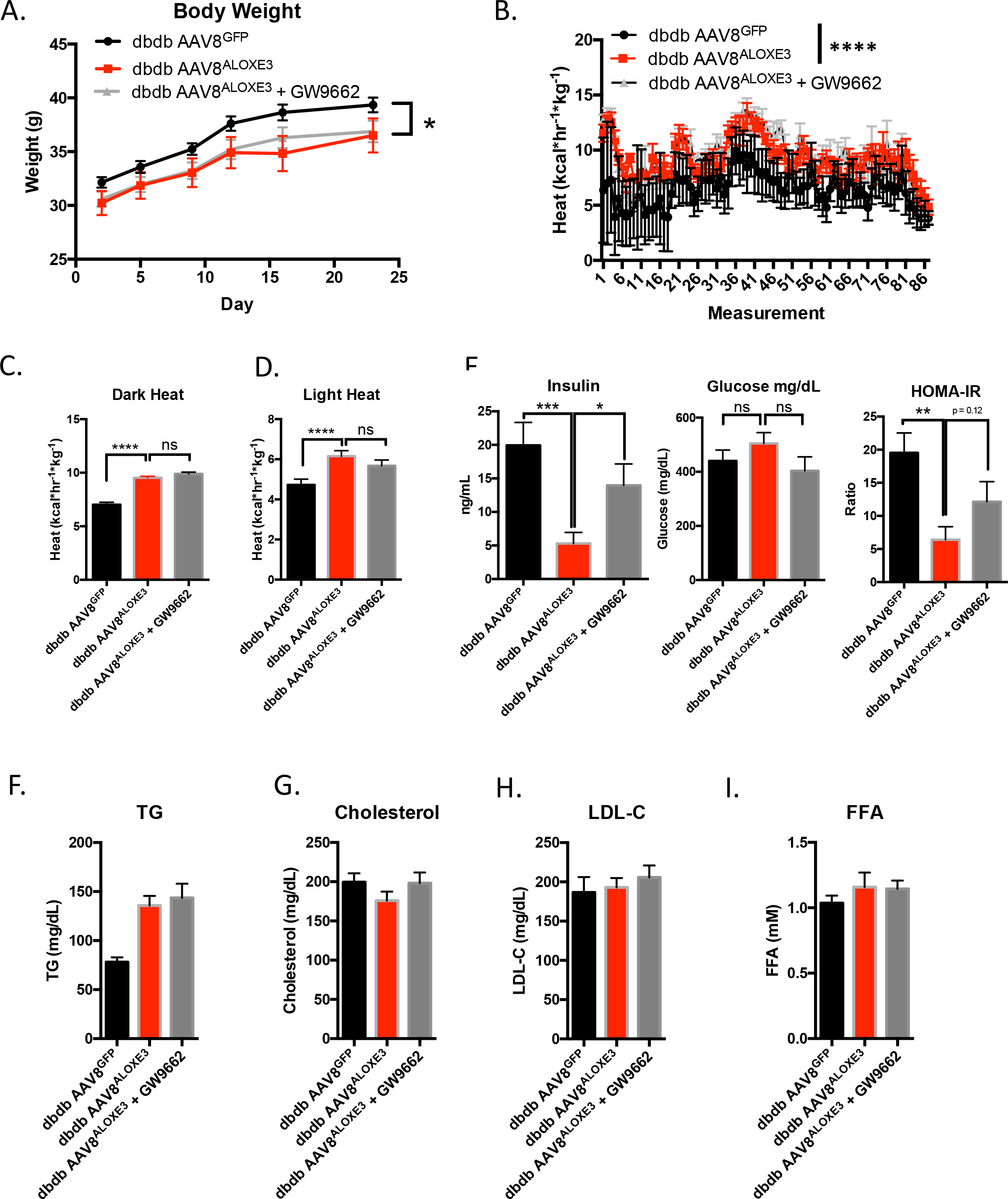
Enhanced whole-body energy metabolism in db/db diabetic mice overexpressing ALOXE3. A. Body weight over time in db/db mice expressing empty vector or ALOXE3 in the presence or absence of GW9662. B. Heat generation over time in db/db mice expressing GFP or ALOXE3. C. and D. Indirect calorimetric quantification of light- and dark-cycle heat generation in db/db mice expressing ALOXE3 or GFP treated with or without GW9662. E. Fasting serum insulin determined by ELISA, serum glucose determined by colorimetric assay, calculated HOMA-IR indcex based on glucose and insulin data. F-I. Serum TG, cholesterol, LDL-C and FFA content in db/db mice with or without hepatic ALOXE3 overexpression and with or without GW9662 treatment. *, ** and ***, P < 0.05, < 0.01, < 0.001 and < 0.0001 between groups. n = 5-10 mice per experiment.

**Figure 9.**
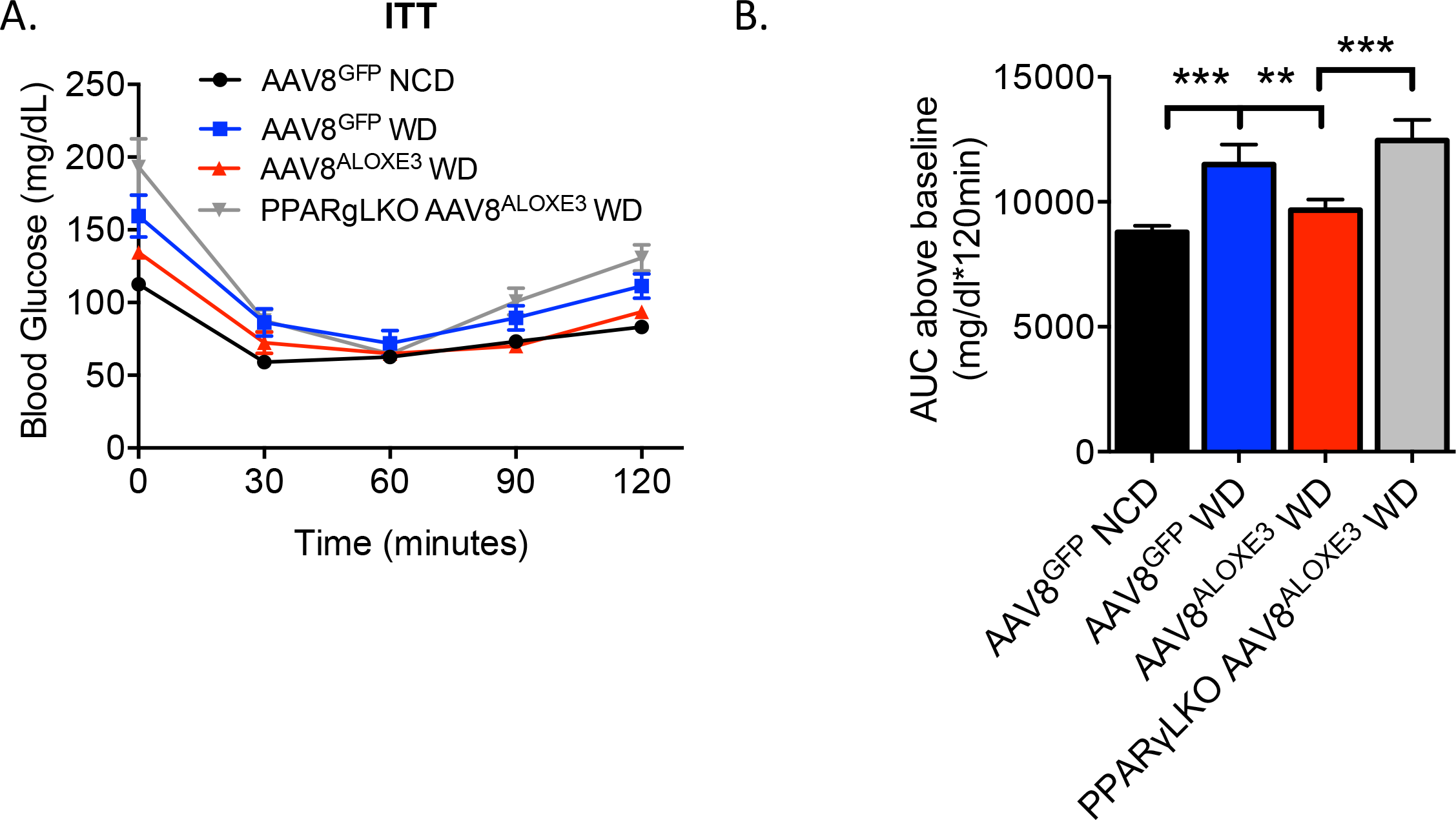
Insulin sensitization by ALOXE3 requires PPARγ. A. Insulin tolerance testing in WT or PPARγ LKO mice expressing hepatic GFP or ALOXE3 after chow or Western diet. B. Quantification of total area under the glucose-time curve. ** and ***, P < 0.01 and 0.001 between groups. n = 4-5 mice per experiment.

Because hepatic ALOXE3 expression enhanced peripheral insulin sensitivity in HTFC-fed mice (Fig. 4), and because PPARγ agonism by TZDs enhances peripheral insulin sensitivity (33,34), we evaluated whether ALOXE3 enhances peripheral insulin and glucose homeostasis dependent on PPARγ and yet independent of the leptin receptor. Fasting circulating insulin and HOMA-IR were significantly lower in ALOXE3^db/db^ mice without changes in circulating glucose when compared with db/db mice expressing GFP. The reduction in circulating insulin and HOMA-IR were reversed in mice concomitantly treated with the PPARγ inhibitor, GW9662, suggesting that ALOXE3 improves glucose and insulin homeostasis in a PPARγ-dependent. In contrast with our diet-induced model, targeted hepatic ALOXE3 expression did not reduce circulating lipids in leptin receptor-deficient mice. Together, the data in our leptin-deficient model elucidate leptin- and PPARγ-dependent functions by ALOXE3.

To gain further specificity regarding the role of hepatocyte PPARγ in ALOXE3-enhanced insulin sensitivity, we generated mice harboring a hepatocyte-specific PPARγ deletion (hereafter, “PPARγ LKO mice”) by crossing mice with homozygous floxed PPARγ alleles with mice expressing cre recombinase driven by the albumin promoter. Mice were placed on a chow or Western diet to induce insulin resistance over 12 weeks, then subjected to insulin tolerance testing. Area under the ITT curve analysis revealed that WD feeding increased AUC in GFP-overexpressing animals, whereas AUC was reduced in ALOXE3-overexpressing animals. In striking contrast, WD-fed ALOXE3-overexpressing PPARγ LKO mice had a significantly elevated AUC when compared with WD-fed ALOXE3-overexpressing mice harboring wild-type hepatic PPARγ.

## DISCUSSION

The concept of the fasting and caloric restriction response as a therapeutic to be leveraged in longevity, healthspan, and metabolism from rodents to early primates to humans (1,35–40) is gaining traction. Already in the clinical realm, caloric restriction improved multiple indices related to aging, cardiovascular disease and diabetes mellitus (2,41–43). However, strict and prolonged adherence to caloric restriction, intermittent fasting or any intensive dietary restriction directed toward weight loss carries with it several practical barriers. Therefore, identifying the therapeutic determinants of the hepatic glucose fasting response opens the promise of novel, targeted and sustainable treatments against metabolic and other diseases. Here, we show that ALOXE3 is activated by the adaptive hepatic fasting response, and is itself sufficient to enhance insulin sensitivity, increase basal caloric expenditure, reduce diet-induced weight gain and fat accumulation, hepatic steatosis and dyslipidemia.

Fully defining the metabolic functions of ALOXE3 has been previously limited by its basal tissue expression distribution and by the dramatic phenotype observed in animals and humans deficient for this enzyme. ALOXE3 was first identified as an epidermal-type lipoxygenase that mediates skin differentiation via hydroperoxide isomerase (26,44). Humans born with a defect in this gene develop ichthyosiform disease, characterized by profound skin barrier dysfunction and transepidermal water loss (27,29). This was confirmed in murine models of ALOXE3 deficiency, which succumb to massive dehydration without intervention shortly after birth (45). Subsequent work demonstrated a role for ALOXE3 in adipocyte differentiation via PPARγ agonism in cultured adipocytes as well (28). This introduced the possibility that ALOXE3 has important functions that extend beyond epidermal tissues. However, a potential function in liver was largely ignored, perhaps in part because non-fasting mRNA is low when compared with epidermis and tongue (45) by Northern Blot detection. However, we show here that serum and glucose withdrawal, trehalose and lactotrehalose (trehalose analog) treatment, and flavone-class (quercetin) GLUT inhibitor treatment in vitro, as well as trehalose feeding and acute fasting in vivo each robustly induce ALOXE3 protein and mRNA expression. This raised the prospect that ALOXE3 mediates part of the physiological hepatic fasting response. We postulate that the observed increased insulin sensitivity in hepatic ALOXE3-overexpressing mice reflects an adaptive response that enhances peripheral insulin sensitivity in preparation for a subsequent meal after prolonged fasting. This function would be particularly useful to maximize macronutrient absorption for animals in which meals are few and far between (e.g. intermittently fasting or hibernating animals). Indeed, this teleological rationale has been proposed for other physiological fasting-induced pathways, such as FGF21 signaling (46,47).

The data herein suggest that ALOXE3 ameliorated the diet-induced obesity and *db/db* diabetic phenotype at least in part via a PPARγ-dependent mechanism - most prominently with regard to the insulin sensitizing effects of ALOXE3. ALOXE3 overexpression and trehalose treatment in hepatocytes induced the known PPARγ ligand, 12-KETE (19), and PPARγ blockade by GW9662 administration in isolated hepatocytes and in vivo reversed ALOXE-mediated changes in hepatocyte oxygen consumption rate, insulin resistance, and PGC1α upregulation. Moreover, liver-specific PPARγ deletion abrogated ALOXE3-enhanced insulin sensitivity. Together, these data implicate a novel hepatic ALOXE3-PPARγ axis that improves whole-body insulin sensitivity. These findings are consistent with the previously demonstrated control of peripheral insulin sensitivity via hepatic PPARγ in ob/ob mice (31,32). However, given that GW9662 did not significantly reverse ALOXE3 effects on hepatic fat, however, we cannot rule out a model in which ALOXE3 exerts both PPARγ-dependent and -independent functions to enhance hepatic and extrahepatic metabolic homeostasis. This model is also supported by the lack of effect of GW9662 also on ALOXE3-induced thermogenesis. In addition, it should be recognized that genetic disruption of both 12-LOX and 15-LOX in multiple diabetic models largely recapitulates the ALOXE3-mediated enhancement of hepatic insulin sensitivity and reduced hepatic steatosis (15–17). Therefore, it is quite feasible that – in addition to generating the PPARγ ligand 12-KETE – either ALOXE3 activity also depletes 12-LOX products 12-HpETE and 12-HETE (Fig. 2) to the host’s benefit, or 12-LOX and 15-LOX deletion shunted arachidonic acid down the ALOXE3 enzymatic pathway to activate PPARγγ.

The current study examines therapeutic effects of hepatic ALOXE3 induction. From a clinical therapeutic perspective, virus-based genetic therapy for many monogenic diseases has already reached clinical care (48). However, this therapeutic approach is not yet fully optimized for polygenic diseases, such as obesity, insulin resistance, and non-alcoholic fatty liver disease (49). Therefore, when compared with precision small-molecule therapeutics, a genetic overexpression approach at present is likely to be a more distant clinical therapy. And although the full extent of ALOXE3 regulation has yet to be elucidated as a means to leverage ALOXE3-enhancing pathways, we demonstrated here that ALOXE3 is potently upregulated by generalized stimuli such as fasting, trehalose feeding, serum withdrawal, and flavone-class GLUT inhibition. In contrast, canonical fasting intermediates: AMPK, PGC1α, FGF21, PPARα were dispensable for trehalose-induced ALOXE3 expression. Accordingly, SIRT1 overexpression was insufficient to upregulate ALOXE3 in vivo. It is therefore intriguing that we were able to achieve enhanced in vitro ALOXE3 upregulation comparable to viral ALOXE3 overexpression by treatment with the trehalase-resistant trehalose analogue, lactotrehalose (22–24,50,51). Our characterization of this glucose-galactose trehalose analogue, and the finding that lactotrehalose exhibits enhanced fasting-mimetic potency is a critical advance for at least two reasons. First, abundant trehalase expression in human gut, liver, brain, kidney and reproductive tissues raises the possibility that trehalose-based therapies may be limited by trehalase-mediated trehalose degradation. By corollary, trehalase resistant compounds such as lactotrehalose are postulated to confer enhanced efficacy. Secondly, it is well understood that GLUT8 and GLUT2 (the two most highly expressed hepatic glucose transporters (7)) are competitively inhibited by galactose (52–54). And in light of the fact that galactose is one of the saccharide moieties that comprises lactotrehalose, the data suggest that altering the saccharide moieties in trehalose modulates potency (and perhaps also selectivity) based on each GLUT’s substrate predilection.

Taken together, although greater detail regarding intermediary regulation of ALOXE3 is warranted, augmenting ALOXE3 expression and other components of the hepatic fasting response is now feasible through fasting itself, through glucose withdrawal, flavone-class or trehalose / analogue-class GLUT inhibition. The palatability, and heavy first-pass enterohepatic kinetics (which minimizes peripheral tissue GLUT side-effects) indeed make trehalose and its analogues especially attractive candidate nutraceuticals to elicit this response. Future directions should interrogate the fasting response-inducing potential, kinetics and efficacy of trehalose and its analogs in mitigating metabolic disease (21,22,55). In addition, characterizing the extent to which ALOXE3 regulation in other tissues is necessary or sufficient to augment the tissue-specific adaptive fasting response. Finally, understanding the full cadre of epoxyalcohols and other lipid products of ALOXE3 enzymatic activity, and their specific roles in the mitigating metabolic disease are of special pharmaceutical and clinical interest (15,56–59).

In summary, we identified the lipoxygenase ALOXE3 as a novel effector of the hepatic fasting response that is sufficient to augment basal caloric expenditure, ameliorate insulin resistance, weight gain and hepatosteatosis. The rapidly rising prevalence of each of these major public health problems throughout the Western and Developing worlds mandates novel therapeutic pathway generation and leveraging thereof. We assert that further interrogation into how hepatic glucose transport mediates the networked adaptive hepatic fasting response will advance the field toward new and effective human therapy against metabolic disease.

## MATERIALS AND METHODS

### Mouse Models and Treatment

All animal procedures were approved by the Washington University School of Medicine Animal Studies Committee. All mice were caged in specific pathogen-free barrier housing with 12h light-dark cycles and free access to water and rodent chow. For transgenic studies, WT C57B/6J mice and *Lepr*^db/db^ mice were obtained directly from the Jackson Laboratory (Bar Harbor, ME). Upon arrival, mice were equilibrated for a minimum of 7 days in the specific pathogen-free vivarium with automated 12h light-dark cycles and free access to water and rodent chow. prior to initiating metabolic measurements. ATG16L1^HM^ mice were a provided by Herbert “Skip” W. Virgin’s laboratory (60). PPARγLKO mice were obtained directly from the Jackson Laboratory (Bar Harbor, ME) to the laboratory of David Rudnick (Washington University) and after propagation were provided for experimentation.

Adeno-associated viruses overexpressing either GFP or ALOXE3 under control of the thyroid binding globulin (TBG) promoter were obtained as ready-to-use viral stocks from Vector Biolabs (Philadelphia, PA). 10^11^ viral particles per animal were injected via tail vein in 6 week-old mice 10 days prior to dietary initiation or 38 days prior to sacrifice in the genetic obesity model (e.g. 10 days rest period +. ALOXE3 expression was quantified in pilot experiments by qRT-PCR analysis of hepatic, skeletal muscle, brown and epididymal white adipose tissues 10 days post-injection and confirmed selective hepatic ALOXE3 overexpression (not shown).

Antisense oligonucleotides (ASOs) were obtained from IONIS Pharmaceuticals as ready-to-inject oligomers, and were used precisely as described previously (61). GW9662 was obtained from Cayman Chemicals (Ann Arbor, MI, #70785).

### Lactotrehalose Synthesis and Purification

Synthesis and purification of lactotrehalose was carried out precisely as described (62). We confirmed 98-99% purity by ^1^H-NMR (not shown).

### Serum analyses

Fasting blood glucose was measured via glucometer using tail vein blood. For all other serum analyses, submandibular blood collection was performed immediately prior to sacrifice and serum was separated. Insulin ELISA (Millipore #EZRMI-13K), Triglycerides (Thermo Fisher Scientific #TR22421), Cholesterol (Thermo Fisher Scientific #TR13421), and Free Fatty Acids (Wako Diagnostics #999-34691, #995-34791, #991-34891, #993-35191) quantification were performed using commercially available reagents according to manufacturer’s directions. Albumin levels were quantified using an AMS LIASYS Chemistry Analyzer.

### Hepatic lipids

Lipids were extracted from ~100 mg hepatic tissue homogenized in 2:1 chloroform:methanol. 0.25% - 0.5% of each extract was evaporated overnight prior to biochemical quantification of triglycerides, LDL-C, cholesterol, and free fatty acids using reagents described above, precisely according to manufacturer’s directions.

### Oil red-O staining

Methanol-fixed frozen sections from WT and transgenic mice were stained according to described protocols (9,11,63).

### Body composition analysis

Body composition analysis was carried out in unanesthetized mice as described (12,63,64) using an EchoMRI 3-1 device (Echo Medical Systems) via the Washington University Diabetic Mouse Models Phenotyping Core Facility. The first 4hrs in the cage were unmeasured in order to allow for acclimation time. Thereafter, the oxygen and CO_2_ consumption and production were quantified for a minimum of one light cycle (0601-1800 hours) and for one dark cycle (1801-0600 hours). VO_2_, VCO_2_, heat, movement and respiratory exchange ratio (RER) were automatically calculated using TSE Phenomaster software.

### In vitro Metabolism (Seahorse XF96) Assays

Seahorse assays were performed using a Seahorse XF96 analyzer Mito Stress Test Kit (Agilent Technologies, Santa Clara, CA) precisely according to manufacturer specifications. Hepatocyte cultures were seeded at 20,000 cells per well prior to assay.

### Immunoblotting

Immunoblotting was performed as described (64). PGC1α antibody was a gift provided by Alan L Schwartz (65), originally generated in the laboratory of Daniel Kelly (66). Antibodies from Cell Signaling Technologies (Beverly MA) were: ²-Actin (#4970); GAPDH (#5174). ALOXE3 (#ab118470) and UCP1 antisera (#ab155117) were from Abcam (Cambridge, MA). LC3B antiserum was obtained from Novus Biologicals (Littleton, CO) (#NB-100-2200).

### Indirect Calorimetry

Oxygen consumption, CO_2_ production, respiratory exchange ratio, and heat production were measured using the Phenomaster system (TSE) via the Washington University Diabetic Mouse Models Phenotyping Core Facility as described (12,64). Metabolic parameters were documented every 13 minutes.

### Cell cultures and treatment

Primary murine hepatocytes obtained from WT mice were isolated as described (9,63) and cultured and maintained in regular DMEM growth media (Sigma, #D5796) containing 10% FBS. For *in vitro* starvation experiments, “starved” media contained 1g/L glucose and 0.5% FBS was used. Cultures were lysed in Trizol and subjected to downstream analysis. *In vitro* genetic knockdown was achieved via siRNA transfection using Lipofectamine 3000 from Invitrogen (L3000015). Trehalose was obtained from Sigma Aldrich (St. Louis, MO) and was ≥ 97% purity by HPLC. 3% trehalose water (w/v) fed ad libitum was used in all in vivo experiments. Baicalein was obtained from Cayman Chemicals (Ann Arbor, MI).

### Quantitative Real-Time RT-PCR (qRT-PCR)

qRT-PCR was performed as previously reported (11,63) with some modifications. Snap-frozen livers or cultured hepatocytes were homogenized in Trizol reagent (Invitrogen #15596026). RNA isolated according to the manufacturer’s protocol was reverse-transcribed using the Qiagen Quantitect reverse transcriptase kit (Qiagen #205310). cDNA was subjected to quantitative PCR using the SYBR Green master mix reagent (Applied Biosystems #4309155). Primer sequences used are available upon request.

### RNAseq

RNAseq was performed by the Washington University Genome Technology Access Center (GTAC). Library preparation was performed with 10ug of total RNA with a Bioanalyzer RIN score greater than 8.0. Ribosomal RNA was removed by poly-A selection using Oligo-dT beads (mRNA Direct kit, Life Technologies). mRNA was then fragmented in buffer containing 40mM Tris Acetate pH 8.2, 100mM Potassium Acetate and 30mM Magnesium Acetate and heating to 94 degrees for 150 seconds. mRNA was reverse transcribed to yield cDNA using SuperScript III RT enzyme (Life Technologies, per manufacturer’s instructions) and random hexamers. A second strand reaction was performed to yield ds-cDNA. cDNA was blunt ended, had an A base added to the 3’ ends, and then had Illumina sequencing adapters ligated to the ends. Ligated fragments were then amplified for 12 cycles using primers incorporating unique index tags. Fragments were sequenced on an Illumina HiSeq-3000 using single reads extending 50 bases.

RNA-seq reads were aligned to the Ensembl release 76 top-level assembly with STAR version 2.0.4b. Gene counts were derived from the number of uniquely aligned unambiguous reads by Subread:featureCount version 1.4.5. Transcript counts were produced by Sailfish version 0.6.3. Sequencing performance was assessed for total number of aligned reads, total number of uniquely aligned reads, genes and transcripts detected, ribosomal fraction known junction saturation and read distribution over known gene models with RSeQC version 2.3.

To enhance the biological interpretation of the large set of transcripts, grouping of genes/transcripts based on functional similarity was achieved using the R/Bioconductor packages GAGE and Pathview. GAGE and Pathview were also used to generate pathway maps on known signaling and metabolism pathways curated by KEGG.

### Liquid chromatography-tandem mass spectral analysis of eicosanoids

Eicosanoids (5-HETE, 12-HETE, and 12-KETE) were extracted with 200 μL of 1:1 methanol/water, containing 2 ng of deuterated 12-HETE-d_8_ (Cayman Chemical, Ann Arbor, MI) as the internal standard. The analysis for eicosanoids as well as the internal standard was performed with a Shimadzu 20AD HPLC system (Columbia, MD) a LeapPAL autosampler (Carrboro, NC) coupled to a tandem mass spectrometer (API 4000, API-Sciex, Foster City, CA) operated with negative ion MRM mode. The sample was injected on to a Thermo-Keystone betasil C-18 HPLC column (2 × 100 mm, 3 μm, Bellefonte, PA) with mobile phases (A: 5 mM ammonium fluoride in water, B: 100 % acetonitrile). The data processing was conducted with Analyst 1.6.3 (API-Sciex).

### Insulin tolerance testing

Insulin tolerance testing was performed precisely as reported previously. Briefly, mice were fasted 4h prior to intraperitoneal administration of 0.75U/kg insulin. Glucose was monitored by glucometer (One Touch Ultra, Lifescan, Wayne, PA) over the course of 2 hours following injection.

### Statistics

Data were analyzed using GraphPad Prism version 6.0 (RRID:SCR_015807). p < 0.05 was defined as statistically significant. Data shown are as mean ± SEM. Unpaired homoscedastic T-tests with Bonferroni *post hoc* correction for multiple comparisons were used for all analyses unless otherwise noted in the Figure Legends. Two-way ANOVA was also used for analyses with two independent variables.

## ACKNOWLEDGEMENTS

This work was supported by the Office of the Assistant Secretary of Defense for Health Affairs, through the Peer Reviewed Medical Research Program under Award No. W81XWH-17-1-0133. Opinions, interpretations, conclusions and recommendations are those of the author and are not necessarily endorsed by the Department of Defense. This work was also supported by grants from the NIH/National Center for Advancing Tranlational Sciences (NCATS) grant #UL1TR002345, the Children’s Discovery Institute (MI-FR-2014-426) AGA-Gilead Sciences Research Scholar Award in Liver Disease, the Washington University Digestive Disease Research Core Center (BJD) (P30DK52574), the Washington University Diabetes Research Center (P30DK020579), Nutrition & Obesity Research Center (P30DK056341), and the Robert Wood Johnson Foundation. BJD is a Scholar of the Washington University Child Health Research Center (K12HD076224), and of the Children’s Discovery Institute (MI-FR-2014-426). BMS was supported by National Institutes of Health (R15 AI117670) to B.M.S. ALM was supported by the Washington University Spencer T. Olin Fellowship, the Washington University NIGMS Institutional Training Grant in Cell and Molecular Biosciences (T32GM007067), and National Science Foundation Graduate Student Fellowship (DGE-1143954). We thank Herbert Virgin at Washington University School of Medicine, for the ATG16L1^HM^ mouse line, and David Rudnick (Washington University) for the liver-specific PPARγ-deficient mice.

## AUTHOR CONTRIBUTIONS

BJD conceived and coordinated the study and wrote the paper. CBH, YZ, ALM, HF, MJB, BMS, AIS and BJD designed, performed and analyzed the experiments. All authors reviewed the results and approved the final version of the manuscript.

## DISCLOSURE OF INTEREST

The authors declare they have no relevant disclosures of interest or financial disclosures.

## REFERENCES

1. Longo VD, Mattson MP. Fasting: Molecular mechanisms and clinical applications. Cell Metab [Internet]. 2014;19(2):181–92. Available from: http://dx.doi.org/10.1016/j.cmet.2013.12.008

2. Barnosky AR, Hoddy KK, Unterman TG, Varady KA. Intermittent fasting vs daily calorie restriction for type 2 diabetes prevention: A review of human findings. Transl Res [Internet]. 2014;164(4):302–11. Available from: http://dx.doi.org/10.1016/j.trsl.2014.05.013

3. Settembre C, De Cegli R, Mansueto G, Saha PK, Vetrini F, Visvikis O, et al. TFEB controls cellular lipid metabolism through a starvation-induced autoregulatory loop. Nat Cell Biol [Internet]. 2013;15(6):647–58. Available from: http://www.ncbi.nlm.nih.gov/pubmed/23604321

4. Nakamura MT, Yudell BE, Loor JJ. Regulation of energy metabolism by long-chain fatty acids. Prog Lipid Res [Internet]. 2014;53(1):124–44. Available from: http://dx.doi.org/10.1016/j.plipres.2013.12.001

5. Chalkiadaki A, Guarente L. Sirtuins mediate mammalian metabolic responses to nutrient availability. Nat Rev Endocrinol [Internet]. 2012;8(5):287–96. Available from: http://dx.doi.org/10.1038/nrendo.2011.225

6. Degirolamo C, Sabbà C, Moschetta A. Therapeutic potential of the endocrine fibroblast growth factors FGF19, FGF21 and FGF23. Nat Rev Drug Discov [Internet]. 2015;15(1):51–69. Available from: http://dx.doi.org/10.1038/nrd.2015.9%5Cn10.1038/nrd.2015.9%5Cnhttp://www.nature.com/doifinder/10.1038/nrd.2015.9

7. DeBosch BJ, Chen Z, Saben JL, Finck BN, Moley KH. Glucose transporter 8 (GLUT8) mediates fructose-induced de Novo lipogenesis and macrosteatosis. J Biol Chem [Internet]. 2014 Apr 18 [cited 2014 Aug 12];289(16):10989–98. Available from: http://www.pubmedcentral.nih.gov/articlerender.fcgi?artid=4036240&tool=pmcentrez&rendertype=abstract

8. DeBosch BJ, Chen Z, Finck BN, Chi M, Moley KH. Glucose Transporter-8 (GLUT8) Mediates Glucose Intolerance and Dyslipidemia in High-Fructose Diet-Fed Male Mice. Mol Endocrinol [Internet]. 2013;27(11):1887–96. Available from: https://academic.oup.com/mend/article-lookup/doi/10.1210/me.2013–1137

9. DeBosch BJ, Heitmeier M, Mayer AL, Higgins CB, Crowley J, Kraft TE, et al. Trehalose inhibits solute carrier 2A (SLC2A) proteins to induce autophagy and prevent hepatic steatosis. Sci Signal. 2016;9(416).

10. DeBosch BJ, Heitmeier MR, Mayer AL, Higgins CB, Crowley JR, Kraft TE, et al. Trehalose inhibits solute carrier 2A (SLC2A) proteins to induce autophagy and prevent hepatic steatosis. Sci Signal [Internet]. 2016 Feb 23;9(416):ra21–ra21. Available from: http://stke.sciencemag.org/content/9/416/ra21.abstract

11. Mayer AL, Higgins CB, Heitmeier MR, Kraft TE, Qian X, Crowley JR, et al. SLC2A8 (GLUT8) is a mammalian trehalose transporter required for trehalose-induced autophagy. Sci Rep [Internet]. 2016 Dec 6;6(December):38586. Available from: http://dx.doi.org/10.1038/srep38586

12. DeBosch BJ, Chen Z, Finck BN, Chi M, Moley KH. Glucose transporter-8 (GLUT8) mediates glucose intolerance and dyslipidemia in high-fructose diet-fed male mice. Mol Endocrinol [Internet]. 2013;27(11):1887–96. Available from: http://www.ncbi.nlm.nih.gov/pubmed/24030250

13. Gorovits N, Cui L, Busik J V., Ranalletta M, De-Mouzon SH, Charron MJ. Regulation of hepatic GLUT8 expression in normal and diabetic models. Endocrinology. 2003;144(5):1703–11.

14. Tsekmekidou XA, Kotsa KD, Tsetsos FS, Didangelos TP, Georgitsi MA, Roumeliotis AK, et al. Assessment of association between lipoxygenase genes variants in elderly Greek population and type 2 diabetes mellitus. Diabetes Vasc Dis Res [Internet]. 2018;147916411875624. Available from: http://journals.sagepub.com/doi/10.1177/1479164118756241

15. Martínez-Clemente M, Ferré N, Titos E, Horrillo R, González-Périz A, Morán-Salvador E, et al. Disruption of the 12/15-lipoxygenase gene (Alox15) protects hyperlipidemic mice from nonalcoholic fatty liver disease. Hepatology. 2010;52(6):1980–91.

16. Samala N, Tersey SA, Chalasani N, Anderson RM, Mirmira RG. Molecular mechanisms of nonalcoholic fatty liver disease: Potential role for 12-lipoxygenase. J Diabetes Complications [Internet]. 2018 Feb 10;31(11):1630–7. Available from: http://dx.doi.org/10.1016/j.jdiacomp.2017.07.014

17. Hernandez-Perez M, Chopra G, Fine J, Conteh AM, Anderson RM, Linnemann AK, et al. Inhibition of 12/15-Lipoxygenase Protects Against β-Cell Oxidative Stress and Glycemic Deterioration in Mouse Models of Type 1 Diabetes. Diabetes [Internet]. 2017 Nov 1;66(11):2875 LP–2887. Available from: http://diabetes.diabetesjournals.org/content/66/11/2875.abstract

18. Cole BK, Morris MA, Grzesik WJ, Leone KA, Nadler JL. Adipose tissue-specific deletion of 12/15-lipoxygenase protects mice from the consequences of a high-fat diet. Mediators Inflamm. 2012;2012.

19. Dozsa A, Dezso B, Toth BI, Bacsi A, Poliska S, Camera E, et al. PPARγ-Mediated and Arachidonic Acid-Dependent Signaling Is Involved in Differentiation and Lipid Production of Human Sebocytes. J Invest Dermatol [Internet]. 2013;134(August). Available from: http://www.ncbi.nlm.nih.gov/pubmed/24129064

20. Hammond VJ, Morgan AH, Lauder S, Thomas CP, Brown S, Freeman BA, et al. Novel keto-phospholipids are generated by monocytes and macrophages, detected in cystic fibrosis, and activate peroxisome proliferator-activated receptor-?? J Biol Chem. 2012;287(50):41651–66.

21. Mardones P, Rubinsztein DC, Hetz C. Mystery solved: Trehalose kickstarts autophagy by blocking glucose transport. 2016;9(416):1–4.

22. Walmagh M, Zhao R, Desmet T. Trehalose analogues: Latest insights in properties and biocatalytic production. Int J Mol Sci. 2015;16(6):13729–45.

23. Chaen H, Nakada T, Mukai N, Nishimoto T, Fukuda S, Sugimoto T, et al. Efficient Enzymatic Synthesis of Disaccharide, alpha-D-Galactosyl alpha-D-Glucoside, by Trehalose Phosphorylase from Thermoanaerobacter brockii. J Appl Glycosci. 2001;48(2):135–7.

24. Kim H-M, Chang Y-K, Ryu S-I, Moon S-G, Lee S-B. Enzymatic synthesis of a galactose-containing trehalose analogue disaccharide by Pyrococcus horikoshii trehalose-synthesizing glycosyltransferase: Inhibitory effects on several disaccharidase activities. J Mol Catal B Enzym [Internet]. 2007;49(1):98–103. Available from: http://www.sciencedirect.com/science/article/pii/S1381117707001695

25. O’Donnell VB, Murphy RC. New families of bioactive oxidized phospholipids generated by immune cells: identification and signaling actions Review article New families of bioactive oxidized phospholipids generated by immune cells: identification and signaling actions. 2012;120(10):1985–92.

26. Yu Z, Schneider C, Boeglin WE, Marnett LJ, Brash AR. The lipoxygenase gene ALOXE3 implicated in skin differentiation encodes a hydroperoxide isomerase. Proc Natl Acad Sci U S A. 2003;100(16):9162–7.

27. Fürstenberger G, Epp N, Eckl KM, Hennies HC, Jørgensen C, Hallenborg P, et al. Role of epidermis-type lipoxygenases for skin barrier function and adipocyte differentiation. Prostaglandins Other Lipid Mediat. 2007;82(1-4):128–34.

28. Hallenborg P, Jørgensen C, Petersen RK, Feddersen S, Araujo P, Markt P, et al. Epidermis-Type Lipoxygenase 3 Regulates Adipocyte Differentiation and Peroxisome Proliferator-Activated Receptor γ Activity Epidermis-Type Lipoxygenase 3 Regulates Adipocyte Differentiation and Peroxisome Proliferator-Activated Receptor Activity †. Mol Cell Biol [Internet]. 2010;30(16):4077–91. Available from: http://www.pubmedcentral.nih.gov/articlerender.fcgi?artid=2916447&tool=pmcentrez&rendertype=abstract

29. Mashima R, Okuyama T. The role of lipoxygenases in pathophysiology; new insights and future perspectives. Redox Biol [Internet]. 2015;6:297–310. Available from: http://dx.doi.org/10.1016/j.redox.2015.08.006

30. Yang SJ, Choi JM, Chae SW, Kim WJ, Park SE, Rhee EJ, et al. Activation of peroxisome proliferator-activated receptor gamma by rosiglitazone increases Sirt6 expression and ameliorates hepatic steatosis in rats. PLoS One. 2011;6(2).

31. Gavrilova O, Haluzik M, Matsusue K, Cutson JJ, Johnson L, Dietz KR, et al. Liver peroxisome proliferator-activated receptor gamma contributes to hepatic steatosis, triglyceride clearance, and regulation of body fat mass. J Biol Chem [Internet]. 2003 Sep 5 [cited 2015 Apr 14];278(36):34268–76. Available from: http://www.jbc.org/content/278/36/34268.long

32. Matsusue K, Haluzik M, Lambert G, Yim SH, Gavrilova O, Ward JM, et al. Liver-specific disruption of PPARγ in leptin-deficient mice improves fatty liver but aggravates diabetic phenotypes. J Clin Invest. 2003;111(5):737–47.

33. Hauner H. The mode of action of thiazolidinediones. Diabetes Metab Res Rev. 2002;18(SUPPL. 2):10–5.

34. He L, Liu X, Wang L, Yang Z. Thiazolidinediones for nonalcoholic steatohepatitis: a meta-analysis of randomized clinical trials. :1–9.

35. Macdonald ANDIA. The effects of fasting on the thermogenic, metabolic and cardiovascular responses to infused adrenaline. 2017;(1995):477–90.

36. Ahmet I, Wan R, Mattson MP, Lakatta EG, Talan M. Cardioprotection by intermittent fasting in rats. Circulation. 2005;112(20):3115–21.

37. Wan R, Camandola S, Mattson MP, Drive S. Intermittent fasting and dietary supplementation with 2-deoxy-. Faseb J [Internet]. 2003;17(1):1133–4. Available from: http://www.ncbi.nlm.nih.gov/entrez/query.fcgi?cmd=Retrieve&db=PubMed&dopt=Citation&list_uids=12709404

38. Colman RJ, Anderson RM, Johnson SC, Kastman EK, Simmons H a, Kemnitz JW, et al. Caloric Restriction Delays Disease Onset and Mortality in Rhesus Monkeys. Science (80-). 2009;325(10 July):201–4.

39. Holmes D. Metabolism: Fasting induces FGF21 in humans. Nat Rev Endocrinol [Internet]. 2016 Jan;12(1):3. Available from: http://dx.doi.org/10.1038/nrendo.2015.202

40. Fazeli PK, Lun M, Kim SM, Bredella MA, Wright S, Zhang Y, et al. FGF21 and the late adaptive response to starvation in humans. J Clin Invest. 2015;125(12):4601–11.

41. Alpert W, Hospital M. Cardiovascular Effects of Intensive Lifestyle Intervention in Type 2 Diabetes. N Engl J Med [Internet]. 2013;369(2):145–54. Available from: http://www.nejm.org/doi/10.1056/NEJMoa1212914

42. Redman LM, Ravussin E. Caloric Restriction in Humans: Impact on Physiological, Psychological, and Behavioral Outcomes. Antioxid Redox Signal [Internet]. 2011 Jan 15;14(2):275–87. Available from: http://www.ncbi.nlm.nih.gov/pmc/articles/PMC3014770/

43. Sathananthan M, Shah M, Edens KL, Grothe KB, Piccinini F, Farrugia LP, et al. Six and 12 Weeks of Caloric Restriction Increases β Cell Function and Lowers Fasting and Postprandial Glucose Concentrations in People with Type 2 Diabetes. J Nutr [Internet]. 2015 Sep 5;145(9):2046–51. Available from: http://www.ncbi.nlm.nih.gov/pmc/articles/PMC4548160/

44. Kinzig A, Heidt M, Fürstenberger C, Marks F, Krieg P. cDNA cloning, genomic structure, and chromosomal localization of a novel murine epidermis-type lipoxygenase. Genomics. 1999;58(2):158–64.

45. Krieg P, Rosenberger S, De Juanes S, Latzko S, Hou J, Dick A, et al. Aloxe3 knockout mice reveal a function of epidermal lipoxygenase-3 as hepoxilin synthase and its pivotal role in barrier formation. J Invest Dermatol [Internet]. 2013;133(1):172–80. Available from: http://dx.doi.org/10.1038/jid.2012.250

46. Lin Z, Tian H, Lam KSL, Lin S, Hoo RCL, Konishi M, et al. Adiponectin mediates the metabolic effects of FGF21 on glucose homeostasis and insulin sensitivity in mice. Cell Metab [Internet]. 2013;17(5):779–89. Available from: http://dx.doi.org/10.1016/j.cmet.2013.04.005

47. Markan KR, Naber MC, Ameka MK, Anderegg MD, Mangelsdorf DJ, Kliewer SA, et al. Circulating FGF21 is liver derived and enhances glucose uptake during refeeding and overfeeding. Diabetes. 2014;63(12):4057–63.

48. Ribeil J-A, Hacein-Bey-Abina S, Payen E, Magnani A, Semeraro M, Magrin E, et al. Gene Therapy in a Patient with Sickle Cell Disease. N Engl J Med [Internet]. 2017;376(9):848–55. Available from: http://www.nejm.org/doi/10.1056/NEJMoa1609677

49. Services A, In AL, Services S, In SL, Specialties M, Types C, et al. Mingming Gao Gene Therapy for Obesity: Progress and Prospects. 2017;

50. Urbanek BL, Wing DC, Haislop KS, Hamel CJ, Kalscheuer R, Woodruff PJ, et al. Chemoenzymatic synthesis of trehalose analogues: Rapid access to chemical probes for investigating mycobacteria. ChemBioChem. 2014;16(17):2066–70.

51. Meints LM, Poston AW, Piligian BF, Olson CD, Badger KS, Woodruff PJ, et al. Rapid One-step Enzymatic Synthesis and All-aqueous Purification of Trehalose Analogues. 2017;(120):e54485. Available from: https://www.jove.com/video/54485

52. Schmidt S, Joost H-G, Schürmann A. GLUT8, the enigmatic intracellular hexose transporter. Am J Physiol Endocrinol Metab. 2009;296(January 2009):E614–8.

53. Leturque A, Brot-Laroche E, Le Gall M. GLUT2 mutations, translocation, and receptor function in diet sugar managing. Am J Physiol Endocrinol Metab. 2009;296(5):E985–92.

54. Colville C a, Seatter MJ, Jess TJ, Gould GW, Thomas HM. Transport Inhibitors. 1993;706:701–6.

55. Rundell SR, Wagar ZL, Meints LM, Olson CD, O’Neill MK, Piligian BF, et al. Deoxyfluoro-d-trehalose (FDTre) analogues as potential PET probes for imaging mycobacterial infection. Org Biomol Chem [Internet]. 2016;14(36):8598–609. Available from: http://xlink.rsc.org/?DOI=C6OB01734G

56. Wahli W, Michalik L. PPARs at the crossroads of lipid signaling and inflammation. Trends Endocrinol Metab [Internet]. 2012;23(7):351–63. Available from: http://dx.doi.org/10.1016/j.tem.2012.05.001

57. Wahli W, Braissant O, Desvergne B. Peroxisome proliferator activated receptors: transcriptional regulator of adipogenesis, lipid metabolism and more…Chem Biol [Internet]. 1995;2(5):261–6. Available from: http://www.sciencedirect.com/science/article/pii/1074552195900454

58. Yu Z, Schneider C, Boeglin WE, Brash AR. Epidermal lipoxygenase products of the hepoxilin pathway selectively activate the nuclear receptor PPAR?? Lipids. 2007;42(6):491–7.

59. Boeglin WE, Kim RB, Brash AR. A 12R-lipoxygenase in human skin: mechanistic evidence, molecular cloning, and expression. Proc Natl Acad Sci U S A [Internet]. 1998;95(12):6744–9. Available from: http://www.pubmedcentral.nih.gov/articlerender.fcgi?artid=22619&tool=pmcentrez&rendertype=abstract%5Cnhttp://www.ncbi.nlm.nih.gov/pubmed/9618483%5Cnhttp://www.pubmedcentral.nih.gov/articlerender.fcgi?artid=PMC22619

60. Cadwell K, Liu JY, Brown SL, Miyoshi H, Loh J, Lennerz JK, et al. A key role for autophagy and the autophagy gene Atg16l1 in mouse and human intestinal Paneth cells. Nature [Internet]. 2008;456(7219):259–63. Available from: http://dx.doi.org/10.1038/nature07416

61. Soufi N, Hall AM, Chen Z, Yoshino J, Collier SL, Mathews JC, et al. Inhibiting monoacylglycerol acyltransferase 1 ameliorates hepatic metabolic abnormalities but not inflammation and injury in mice. J Biol Chem. 2014;289(43):30177–88.

62. Bassily RW, El-Sokkary RI, Silwanis BA, Nematalla AS, Nashed MA. An improved synthesis of 4-azido-4-deoxy- and 4-amino-4-deoxy-[alpha],[alpha]-trehalose and their epimers. Carbohydr Res [Internet]. 1993;239:197–207. Available from: http://www.sciencedirect.com/science/article/B6TFF-42V6T69-2K/2/834911c18d91dd9654da76eec3046443

63. DeBosch BJ, Chen Z, Saben JL, Finck BN, Moley KH. Glucose Transporter 8 (GLUT8) mediates fructose-induced de novo lipogenesis and macrosteatosis. J Biol Chem. 2014 Feb;8.

64. DeBosch BJ, Kluth O, Fujiwara H, Schürmann A, Moley K. Early-onset metabolic syndrome in mice lacking the intestinal uric acid transporter SLC2A9. Nat Commun [Internet]. 2014;5:4642. Available from: http://www.ncbi.nlm.nih.gov/pubmed/25100214

65. Trausch-Azar J, Leone TC, Kelly DP, Schwartz AL. Ubiquitin proteasome-dependent degradation of the transcriptional coactivator PGC-1?? via the N-terminal pathway. J Biol Chem. 2010;285(51):40192–200.

66. Leone TC, Lehman JJ, Finck BN, Schaeffer PJ, Wende AR, Boudina S, et al. PGC-1?? deficiency causes multi-system energy metabolic derangements: Muscle dysfunction, abnormal weight control and hepatic steatosis. PLoS Biol. 2005;3(4):0672–87.

